# Investigations into an overlooked early component of painful nociceptive withdrawal reflex responses in humans

**DOI:** 10.1101/2022.11.29.518364

**Authors:** Oumie Thorell, Johannes Ydrefors, Mats Svantesson, Björn Gerdle, Håkan Olausson, David A. Mahns, Saad S. Nagi

## Abstract

**Introduction:** The role of pain as a warning system necessitates a rapid transmission of information from the periphery for the execution of appropriate motor responses. The nociceptive withdrawal reflex (NWR) is a physiological response to protect the limb from a painful stimulus and is often considered an objective measure of spinal nociceptive excitability. The NWR is commonly defined by its latency in the presumed Aδ-fiber range consistent with the canonical view that “fast pain” is signaled by Aδ nociceptors. We recently demonstrated that human skin is equipped with ultrafast (Aβ range) nociceptors. Here, we investigated the short-latency component of the reflex and explored the relationship between reflex latency and pain perception.

**Methods:** We revisited our earlier work on NWR measurements in which, following convention, only those reflex responses were selected that were in the presumed Aδ range (taken to be latencies ≥90 ms in that study). In our current analysis, we expanded the time window to search for shorter latency responses and compared those with pain ratings.

**Results:** In both cohorts, we found an abundance of recordings with short-latency reflex responses. In nearly 90% of successful recordings, only single reflex responses (not dual) were seen which allowed us to compare pain ratings to reflex latencies. We found that shorter latency reflexes were just as painful as those in the conventional latency range.

**Discussion:** We found a preponderance of short-latency painful reflex responses. Based on this finding, we suggest that short-latency responses must be considered in future studies. We predict these might be signaled by the ultrafast nociceptors, warranting further investigation.

## 1 Introduction

The nociceptive withdrawal reflex (NWR) is a physiological response of the limb away from a painful stimulus. It has been investigated both as a tool to probe spinal nociceptive excitability and because of its close association with subjective pain thresholds (de Willer 1977, Chan and Dallaire 1989, Guieu et al. 1992, Sandrini, Arrigo et al. 1993, Biurrun Manresa et al. 2011, Lim et al. 2011, Lim et al. 2012). The NWR response is measured by electromyography and considered to have two latency components: the first component, referred to as RII, is mediated by Aβ or Group II fibers; the second component, referred to as RIII, is mediated by Aδ or Group III fibers (Hugon 1973). The conventional view is that the first or short-latency response is exclusively tactile, and the second or long-latency response is nociceptive (but also see Willer et al. 1978). While the RII-RIII latency cut-off varies across studies, the exclusion of short-latency responses is common practice (de Willer 1977, Dowman 1991, Dowman 1992, Rhudy and France 2007).

Using microneurography, we recently showed that humans, akin to other mammals, are equipped with ultrafast (Aβ range) nociceptors in the skin (Nagi et al. 2019). Considering this finding, we revisited our earlier work on NWR measurements (Ydrefors et al. 2020): in that study, following convention, NWR responses were only selected if they occurred ≥90 ms, consistent with the presumed Aδ-fiber range. Here we expanded the time window to search for shorter latency responses with the hypothesis that those are nociceptive, corresponding to painful sensations.

We found an abundance of short-latency reflex responses, and these were just as painful as those in the conventional latency range, suggesting that by discarding shorter latencies, we may be overlooking valuable quantitative measures of pain processing.

## 2 Material and Methods

### 2.1 Participants

NWR responses and pain ratings were successfully extracted for 20 fibromyalgia patients (FM: age range, all female) and 10 healthy controls (HC: age range, all female). For details on patient eligibility criteria, refer to Ydrefors et al. (2020). Raw data were unavailable for 10 HC and therefore another 10 HC were recruited (18-30 years, all female). The new participants were not age-matched: these data were collected during the pandemic, and it was considered an unnecessary risk to recruit older participants. Additional data collection was approved by the Swedish Ethical Review Authority (Dnr: 2020-04207), and the study procedures complied with the revised Declaration of Helsinki. All participants gave their written informed consent.

### 2.2 Testing procedure and NWR determination

For full details on the testing procedure, refer to Ydrefors et al. (2020). Briefly, electrical stimuli were delivered to the surface of the foot sole using a constant current stimulator generating a train of 5 square wave pulses (1 ms, 200 Hz), and electromyographic (EMG) responses were recorded from the ipsilateral tibialis anterior muscle. In Ydrefors et al. (2020), reflex responses with Z-scores ≥12 were detected using an automated approach. The maximum amplitude (peak amplitude) in the time window of 90 to 150 ms after stimulus onset and the mean amplitude in the -65 to -5 ms pre-stimulus onset (baseline) were determined on a trial-to-trial basis. To determine the Z-score, the difference between peak amplitude and mean baseline amplitude was divided by the baseline standard deviation.

All participants rated the intensity of the sensation, immediately after receiving the electrical stimulus, on a descriptive numeric scale from 0 to 10. Zero corresponded to “no feeling”, 1 to a “slight feeling”, 2 to a “distinct feeling”, 3 to “unpleasantness”, 4 to “just noticeable pain”, 5 to “slight pain”, 6 to “distinct pain”, 7 to “moderately intense pain”, 8 to “intense pain”, 9 to “very intense pain” and 10 to the “worst imaginable pain” (Ydrefors et al. 2020).

### 2.3 New data analysis

The data were pseudonymized and information on latency, Z-scores, NWR thresholds, and age were stored in a relational database. NWR responses were converted from text files (.txt) into graphs (.png), using a script made in Python Distribution (v3.7.4, Python Software Foundation, Beaverton, USA) and latencies were visually inspected by the author (OT) in LabChart (v8.1.16 ADInstruments, Dunedin, New Zealand) and in MATLAB (r2021b, MathWorks Inc, Natick, Massachusetts). Z-scores were calculated for the early time window of 50 to 89 ms after stimulus onset, using the same MATLAB algorithm that was used for the 90 to 150 ms time window. Careful visual inspection of the data allowed us to extract reflex responses with Z-scores ≥6.

### 2.4 Statistical analysis

Statistical analysis was done in GraphPad Prism (v. 9.1.2, GraphPad Software, San Diego, USA). QQ plots, means, standard deviations, and skewness were assessed to determine the normal distribution of the data. Where possible, parametric tests were used. If assumptions for parametric tests were violated, non-parametric tests were conducted.

To compare independent differences between HC and FM, unpaired two-tailed t-test was used, while small sample size data were analysed using two-tailed Mann-Whitney U test. Two-way ANOVA was used to compare multiple independent groups with Tukey’s test as a multiple comparison (post-hoc) test. Only main effects were analyzed due to uneven sample sizes. A statistical value of p < 0.05 was considered statistically significant.

Effect sizes were calculated as Hedges’ g for the unpaired t-tests, due to different sample sizes, and partial eta square (η^2^_p_) for the two-way ANOVA. Common language effect sizes (CLES) are shown for statistically significant ANOVA results. When using non-parametric tests, CLES is shown to compare effects to the parametric results. Effect sizes were calculated in statistical calculators (Lakens 2013, Lenhard 2016). Numbers are presented as mean and standard deviation for parametric tests and median and interquartile range for non-parametric tests.

## 3 Results

Three hundred and eighty-two painful reflex responses with Z-scores ≥6 were successfully extracted from 340 EMG recordings: 166 NWR responses in 20 HC and 216 NWR responses in 20 FM (Fig. 1A-B). The Z-scores ranged from 6.1 to 726.4 with rectified amplitudes of 6 to 698 mV. These reflex responses corresponded to ratings of 4 (“just noticeable pain”) and higher. 63 reflex responses (14.2% of total (382 + 63)) corresponded to ratings below 4 (i.e., nonpainful).

**Figure 1.**
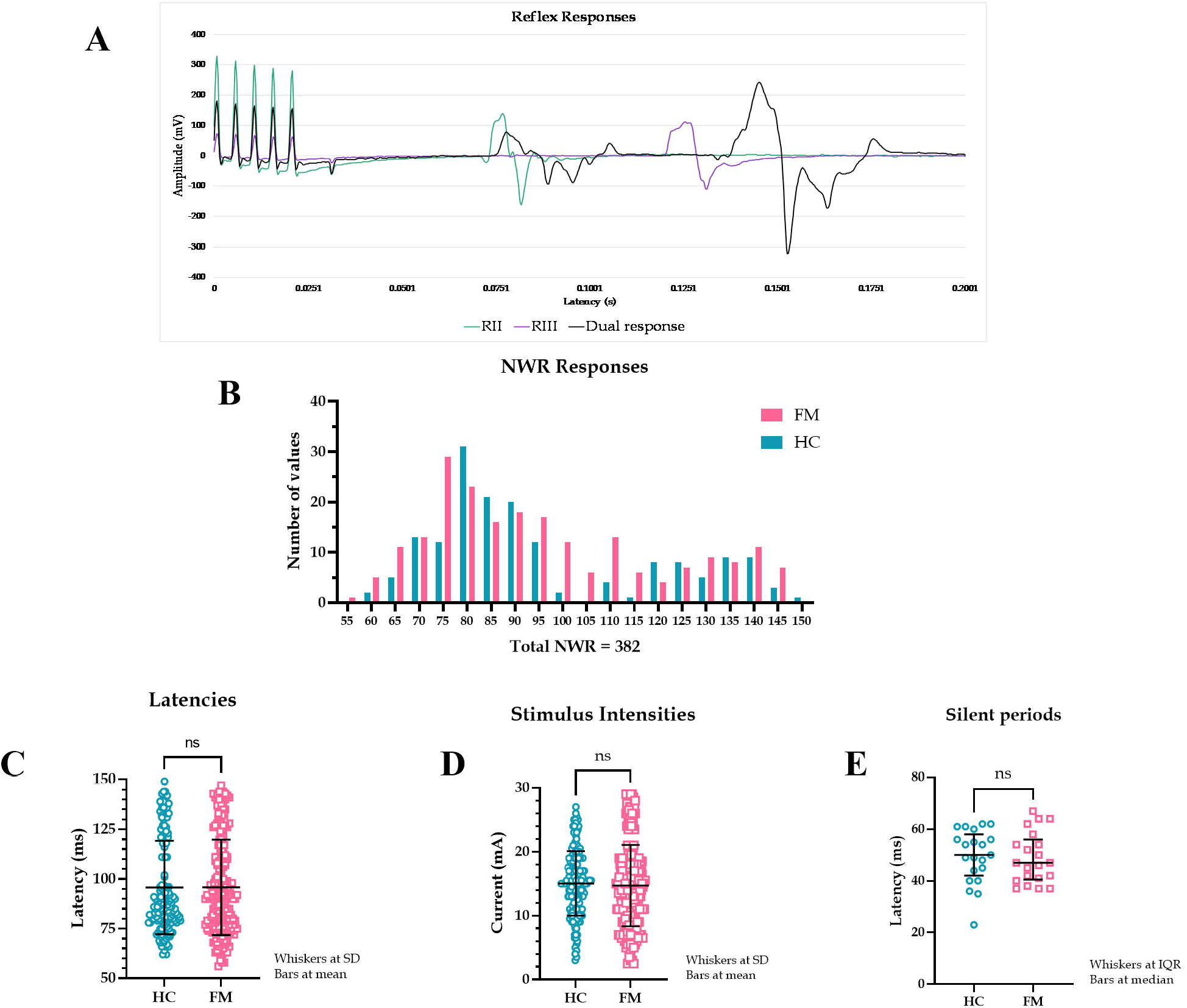
**A**. Examples of reflex recordings with RII, RIII, and dual responses superimposed. The five peaks at the beginning of the graph represent the electrical stimulus (5 square pulses). **B**. Latency spread of painful NWR responses. Healthy controls (HC) had a bimodal distribution while fibromyalgia patients (FM) had a more even distribution throughout the time analysis window. The y-axis shows the number of NWR responses, and the x-axis shows reflex latencies. **C**. Latencies of all NWR responses from HC and FM. Latencies did not differ between the two groups (HC: 95.7 ± 23.5 ms, FM: 95.7 ± 24.0 ms, t(380) = 0.043, p = 0.965, 95% CI [-4.722, 4.936], Hedges’ g = 0.004, CLES = 50.1%). **D**. Stimulus intensities of all NWR responses in HC and FM. Stimulus intensities were not different between HC and FM (HC: 15.0 ± 5.0 mA, FM: 14.7 ± 6.4 mA, t(380) = 0.568, p = 0.570, 95% CI [-1.524, .840], Hedges’ g = 0.059, CLES = 51.7%). **E**. Duration of silent EMG period intervening a dual reflex response occasionally seen. No statistical difference was found in the duration of the silent period between HC and FM (HC: 50.0 (16.0) ms, FM: 47.0 (15.6) ms, U = 203, p = 0.667, CLES = 52.1%).

Reflex latencies and stimulus intensities did not differ between HC and FM groups (Fig. 1C-D). In 42 out of 340 EMG recordings (21 each in HC and FM groups, 12.4% of the total), two reflex responses were seen (84 reflex responses). These dual responses were separated by a silent EMG period (SP) with a duration of 23 to 67 ms (mean 49.1 ms). The SP duration was not different between HC and FM groups (Fig. 1E).

In terms of RII-III prevalence, 192 (50.3%) reflex responses were identified in the 90-to 150-ms latency range corresponding to RIII, and 190 reflexes (49.7%) were identified in the 50-to 90-ms latency range corresponding to RII. The RII data are new: the pre-set 90-ms latency cut-off implemented in Ydrefors et al. (2020) resulted in an automatic discounting of shorter latency responses. To compare pain with NWR responses, only those reflex responses that were painful (at least a 4 rating on a 0-10 scale) were included in the main analysis.

### 3.1 Comparison of RII and RIII responses between FM and HC groups

Eighteen out of 20 HC (90%) and 13 out of 20 FM (65%) had an RII. No differences were found in stimulus intensities between RII and RIII (p = .717) (Fig. 2A). To determine the relationship between subjective pain rating and reflex latency, only single reflex responses were considered (298 reflex responses) (Fig. 2B). FM had higher pain ratings than HC for both RII and RIII responses. Within each group (FM/HC), when pain ratings were compared between RII and RIII, these were not different, suggesting that RII was just as painful as RIII (Fig. 2C). A small proportion of the NWR responses were non-painful: 58 in the HC group and 5 in the FM group (Fig. 2D).

**Figure 2.**
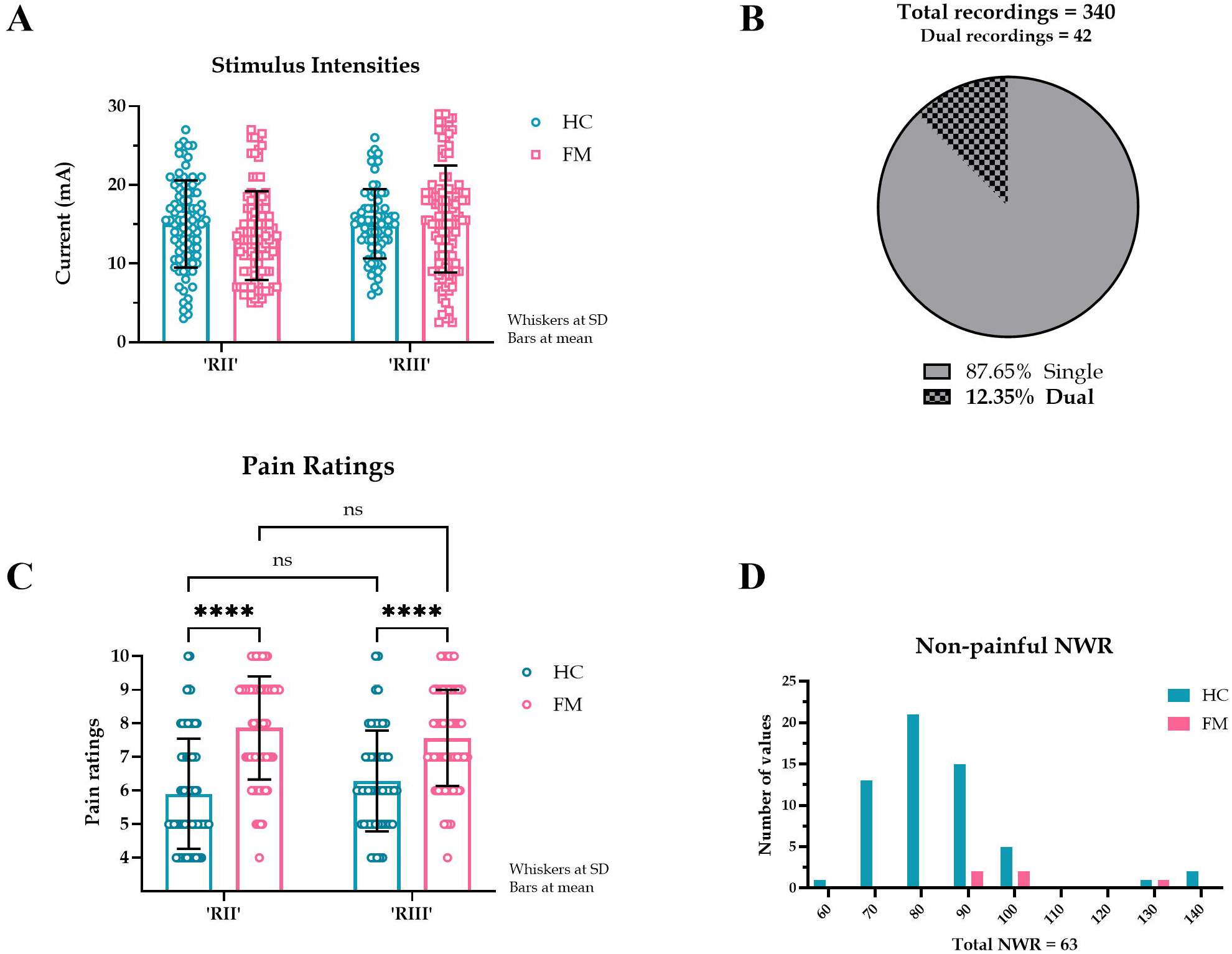
**A**. Stimulus intensities required to evoke RII and RII in HC and FM. A pre-set cut-off of 90 ms was implemented to separate RII and RIII responses. The stimulus intensities required to evoke RII and RIII responses were not different (F(1, 283) = 0.131, p = 0.717, CI [-0.292, 0.424], η2p < 0.000, CLES = 50.0%). There was a main effect of subject type (HC or FM) (F(1, 283) = 79.9, p < 0.000, η2p = 0.022, CLES = 77.3%) but post hoc test indicated no differences in stimulus intensities for subject or reflex type. **B**. Proportion of single and dual NWR EMG recordings. The dual recordings (84 reflex responses) were excluded from subsequent perception analysis. **C**. Pain ratings corresponding to RII and RIII response in HC and FM. Simple main effects indicated that the reflex type had no effect on pain ratings (F(1, 295) = 0.011, p = 0.916, CI [-0.333, 0.370], η2p < 0.001, CLES = 50.0%). Subject type (HC or FM) did have a large effect on pain ratings (F(1, 295) = 82.6, p < 0.001, CI [-2.001, -1.288], η2p = 0.218, CLES = 77.2%). **D**. Latency spread of non-painful NWR responses. Only a few NWR responses (14.6%) were reported as non-painful (pain rating <4), almost entirely by HC (HC: 82 (15.5) ms, n = 58. FM: 96 (25) ms, n = 5). These responses were not included in our analysis. The y-axis shows the number of NWR responses, and the x-axis shows reflex latencies.

As an alternative to a pre-set latency cut-off for RII/III (90 ms), we used the data from dual reflex responses to distinguish between RII and RIII latencies. No dual responses occurred before 93 ms or after 99 ms, therefore we took an in-between value of 96 ms to separate RII and RIII responses (Fig. 3A). Predictably, this increased the proportion of RII responses: in the HC group, the increase was 13.3% (22 additional responses) and in the FM group, the increase was 13.9% (an additional 30 responses) (Fig. 3B-C). However, the overall results did not change (Fig. 3D-E).

**Figure 3.**
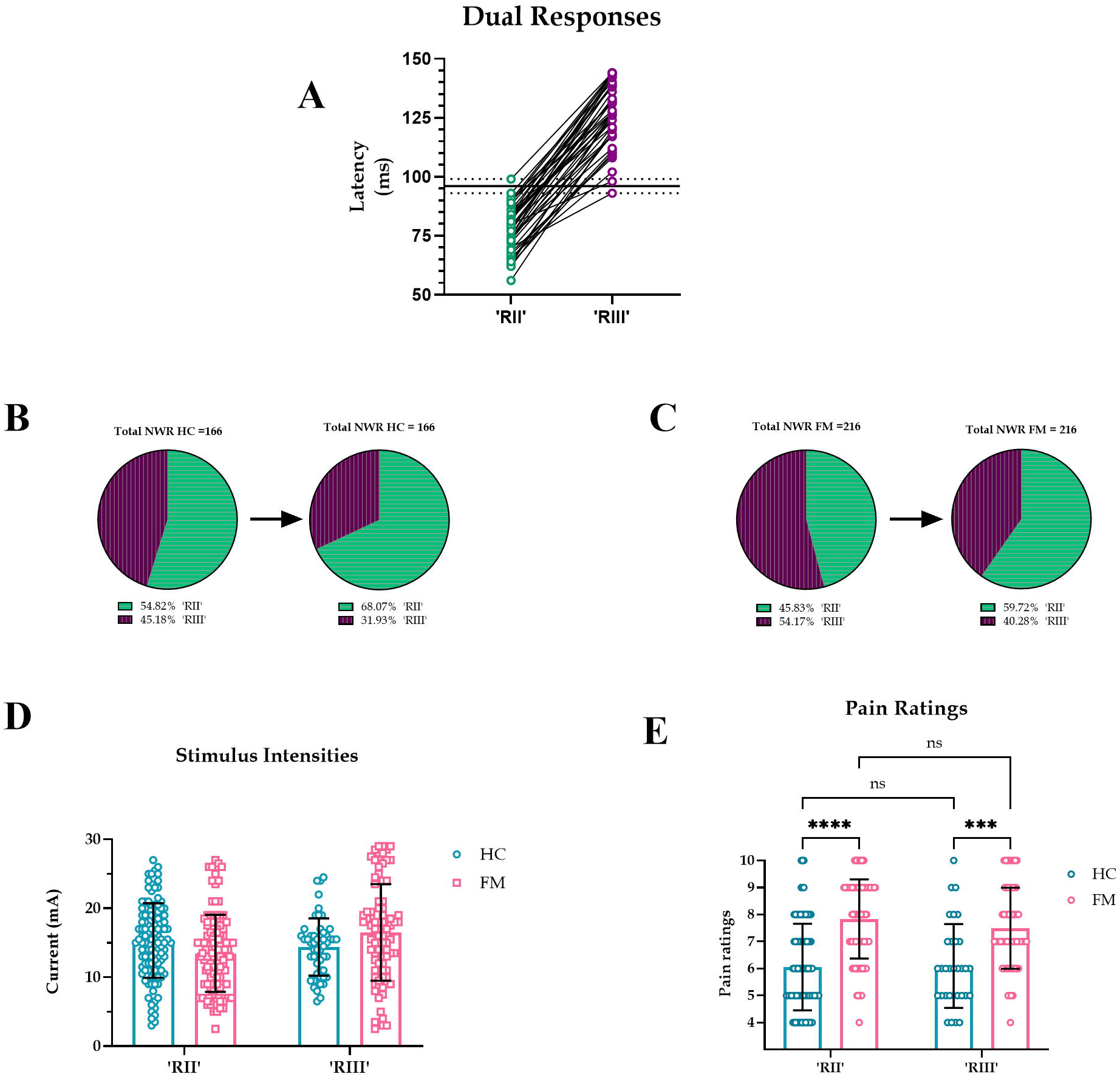
**A**. Latencies of all dual reflex responses. No dual responses were observed before 93 or after 99 ms (dotted lines) so an in-between value of 96 ms was taken to separate RII and RIII responses (solid black line). **B-C**. Proportion of RII and RIII with a divider set at 96 ms. The pie chart on the left shows NWR responses with a pre-set cut-off at 90 ms, and the pie chart on the right shows NWR responses with a divider set at 96 ms, a cut-off derived from dual responses. Predictably, in both HC and FM groups, the prevalence of RII increased and the prevalence of RIII decreased when a 96-ms cutoff was implemented. **D-E**. Stimulus intensities and pain ratings for RII and RIII in HC and FM using 96 ms as the divider. Current intensities had an effect on reflex type (F(1, 379) = 5.262, p = 0.022, CI [-2.630, -0.202], η2p = 0.013, CLES = 56.5%), but not on subject type (F(1,379) = 0.587, p x0025;). Post hoc test indicated no difference in stimulus intensities for subject or reflex type. Subject type had an effect on pain ratings (F(1, 295) = 84.8, p < 0.001, CI [-2.022, -1.310], η2p = .223, CLES = 77.6%) but not reflex type (F(1, 295) = 1.125, p = 0.290, CI [-0.173, 0.577], η2p = .003, CLES = 53.1%).

### 3.2 Z-scores

In Ydrefors et al. (2020), an automated method was used to detect reflex responses with Z-scores of ≥12. Here, visual inspection of the data allowed us to include responses with Z-scores ≥6. For comparison, we performed the analysis implementing the original Z-score (≥12) condition. A total of 234 NWR responses had a Z-score of 12 or higher (Fig. S1A). The proportion of dual responses increased from 12.4% to 17.6% (Fig. S1B). Reflex latencies were shorter in the FM group compared to the HC group (Fig. S1C). Stimulus intensities did not differ between HC and FM (Fig. S1D). FM had higher pain ratings than HC, but pain ratings and stimulus intensities were not different between RII and RIII for either group (Fig. S1E-F).

## 4 Discussion

In this study, we reanalyzed the reflex data from Ydrefors et al. (2020): expanding the time window revealed an abundance of RII NWR responses at stimulus intensities deemed painful. Remarkably, most recordings contained a single reflex response, contrary to the notion that the reflex usually consists of a double burst of EMG activity. We excluded the dual responses from the perception analysis and compared reflex latency (all single responses) with corresponding pain ratings. We found that RII responses were just as painful as RIII responses. The canonical view is that the short-latency component of the NWR response is purely tactile (i.e., nonpainful) and signaled by Aβ low-threshold mechanoreceptors. However, our data show a preponderance of painful reflexes with short latencies.

In the literature, the proposed RII-RIII latency cut-off can be anywhere between 60 and 115 ms post-stimulus onset (for references, see Andersen 2007). Here, we compared data based on a pre-set latency cut-off of 90 ms with a post hoc approach whereby we used dual responses to set the RII-RIII cut-off; this did not change the overall results. In the early reflex work showing double-burst EMG activity, the first EMG response had a lower electrical threshold (Hagbarth 1960, Shahani and Young 1971) and was less painful or non-painful (Hugon 1973, de Willer 1977). In our study, nonpainful reflex responses were rarely observed. Using dual nerve stimulation, Willer and colleagues (1978) were able to evoke an NWR response while the small-fiber inputs were blocked by anesthesia. Indeed, in our single-unit microneurography study, we had confirmed that Aβ nociceptive fibers not only respond to and encode nociceptive stimuli, but also evoke a painful percept when selectively activated during intraneural electrical stimulation (Nagi et al. 2019).

In the FM group, pain ratings were higher and nearly all reflex responses were painful. We observed no latency differences between HC and FM groups except when only responses with Z-scores ≥12 were considered; in that case, FM had shorter latencies than HC. Higher pain ratings and shorter latencies could be attributed to peripheral and/or central sensitization in the patient population (Boureau et al. 1991). We did not detect differences in the SP duration between the two groups. The SPs arise due to postsynaptic inhibition in the motor neurons following a strong electrical stimulus of a muscle or cutaneous nerve. Prolonged SP duration has been previously observed in FM patients (Baek et al. 2016) and is thought to reflect spinal dysregulation.

In Ydrefors et al. (2020), a Z-score of ≥12 was considered a successful muscle response. In that study, a fully automated method was used for NWR detection, therefore the Z-score had to be large enough to ensure that noise would not be interpreted as muscle response. In the current study, we visually inspected all data and were able to reliably detect responses with Z-scores ≥ 6, increasing our sample size by over a third. The conclusions were the same regardless of the Z-score threshold.

### 4.1 Limitations

Raw data were not available from 10 HC in the original sample, therefore new participants had to be recruited. The recruitment was done during the pandemic, and it was considered an unnecessary risk to recruit older participants, thus the new sample is not age matched. Other than that, care was taken to ensure that the experimental protocol was as similar as possible to the original study. A comparison between the two HC samples did not reveal any differences in reflex thresholds or pain ratings.

We did not calculate the conduction velocity (CV) of the NWR. This is difficult to do and involves several variables and assumptions, hence the CV estimations in the literature range anywhere from slow to very fast myelinated afferents. One study based on afferent CVs from single painful shocks with near-nerve stimulation of the tibial nerve reported velocities of 18.5 ± 1.3 m/s with onset latencies between 100-200 ms (Ertekin et al. 1975). Another study based on a train of 5 pulses reported conduction velocities of 49 ± 11 m/s with onset latencies between of 50-100 ms (Ellrich et al. 1998). Classification based on conduction velocity into Aβ and Aδ groups is not clear-cut in humans. In animal studies, the D-hair units are considered a benchmark for the Aδ velocity range (Djouhri and Lawson 2004), however, in humans no detailed account of D-hair units exists.

In conclusion, we found a great many short-latency NWR responses that were as prevalent and as painful as the conventional longer latency NWR responses. Reflex responses that were not painful rarely occurred. Only a minority of NWR recordings consisted of two reflexes. Pain ratings were similar across all latencies, suggesting that the short-latency component is not tactile but nociceptive. We predict this fast signaling involves Aβ nociceptors, warranting further investigation.

## 6 Author Contributions

OT, DM, SSN and HO contributed to the design and conception of this study. MS optimized data analysis. OT wrote the first draft of the manuscript. DM, SSN, HO, JY, BG contributed to subsequent revisions of the manuscript. All authors approved the submitted version.

## 8 Acknowledgments

We thank Magnus Kronander for his helpful contribution to data analysis.

## 10 Supplementary Material

See separate document: “Supplementary Material”

## 11 Data Availability Statement

Raw data will be made available upon request.

