## Supplementary Figure Legend for "Investigations into an overlooked early component of painful nociceptive withdrawal reflex responses in humans"

**Supplementary Figure 1. A.** Latency spread of NWR responses with Z-scores of ≥12. HC showed a bimodal distribution while FM had a more even distribution throughout the time analysis window. The y-axis shows the number of NWR responses, and the x-axis shows the reflex latencies. **B.** Proportion of single and dual NWR EMG recordings with Z-scores ≥12. Most of the NWR recordings comprised a single reflex. Compared to reflex data with Z-scores ≥6, here the proportion of dual responses increased from 12.4% to 17.6%. **C.** Reflex latencies of all NWR responses with Z-scores ≥12. FM had shorter latencies than HC (HC: 105.3 ± 46.5 ms, FM: 98.0 ± 33.8 ms, $t(232) = 2.294$, $p = 0.023$, 95% CI [-13.75, -1.042], Hedges' g = 0.311, CLES = 58.6%). **D.** Stimulus intensities of all NWR responses with Z-scores ≥12. Stimulus intensities were not different between HC and FM (HC: 14.4 ± 5.0 mA, FM: 14.9 ± 6.4 mA, $t(232) = 0.556$, $p = 0.579$, 95% CI [-1.151, 2.055], Hedges' g = .076, CLES = 52.2%). **E.** Pain ratings (≥4) corresponding to RII and RIII responses with Z-scores ≥12. Dual responses were excluded from perception analysis. Simple main effects indicated that the reflex type (RII or RIII) had no effect  on pain ratings ($F(1, 190) = 0.817$, $p = 0.367$, CI [-0.652, 0.242], $\eta2p < 0.004$, CLES = 53.6%), whereas the subject type (HC or FM) had a large effect on pain ratings ($F(1, 190) = 66.03$, $p < 0.001$, CI [-2.287, -1.394], $\eta2p = 0.258$, CLES = 79.8%). **F.** Stimulus intensities required to evoke RII and RII with Z-scores of  ≥12. The subject type had no effect on stimulus intensities ($F(1, 230) = 0.161$, $p < 0.689$, $\eta2p = 0.000$, CLES = 50.0%). The reflex type showed a minor effect on stimulus intensities ($F(1,230) = 4.845$, $p = 0.0287$, CI [-3.568, .197], $\eta2p < 0.021$, CLES = 58.2%) but post hoc test indicated no differences in stimulus intensities between subject or reflex type.
