## Supplementary figures and images for "Investigations into an overlooked early component of painful nociceptive withdrawal reflex responses in humans"

### Supplementary Figure 1.

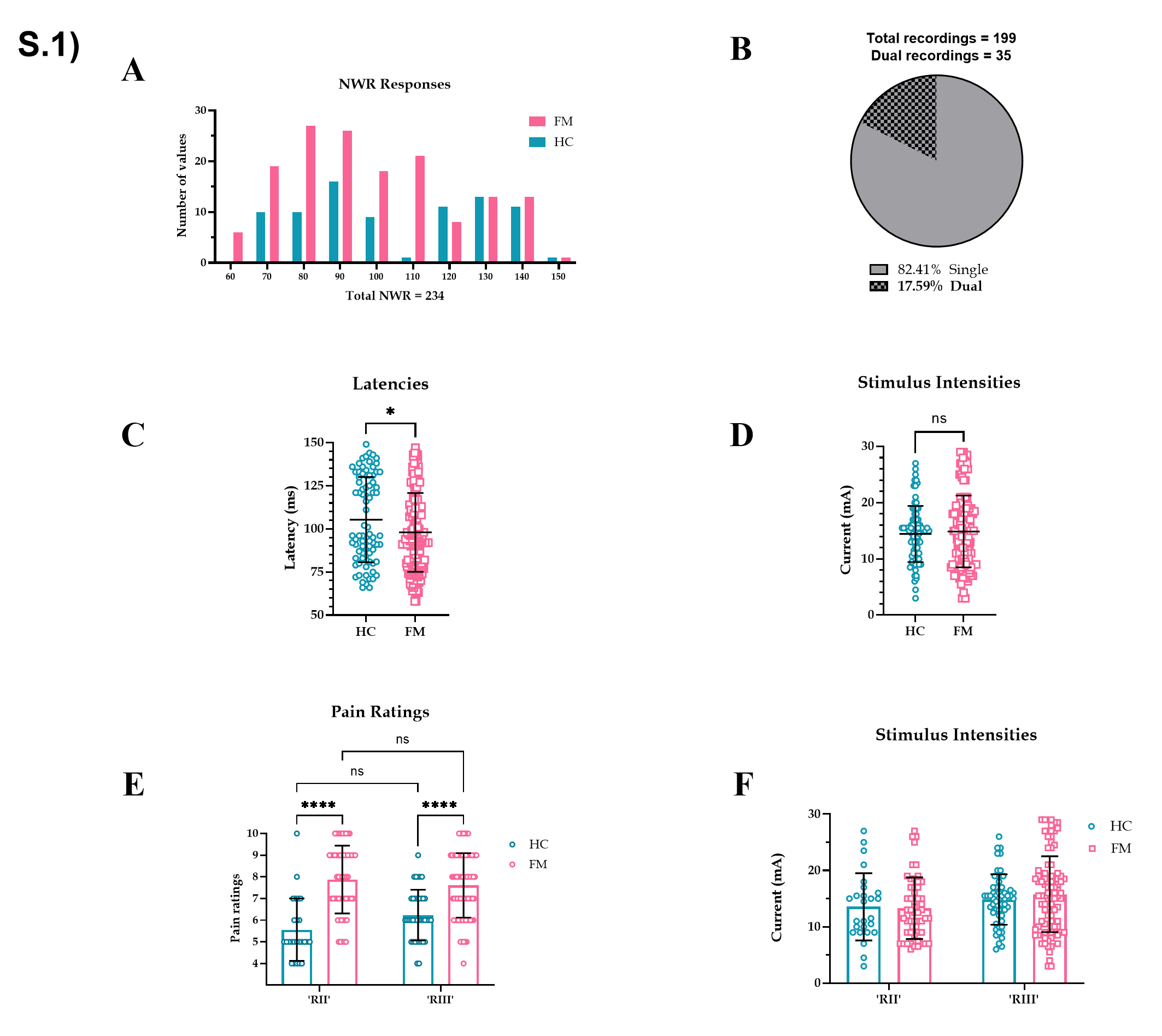
